# An improved method for fitting gamma distribution to substitution rate variation among sites

**DOI:** 10.1101/276329

**Authors:** Xuhua Xia

**Affiliations:** University of Ottawa, Ottawa, Ontario K1N 6N5 Canada

**Keywords:** phylogenetics, rate heterogeneity over sites, substitution model, gamma distribution, simultaneous estimation

## Abstract

Gamma distribution has been used to fit substitution rate variation over site. One simple method to estimate the shape parameter of the gamma distribution is to 1) reconstruct a phylogenetic tree and the ancestral states of internal nodes, 2) perform pairwise comparison between nodes on each side of each branch to count the number of “observed” substitutions for each site, and apply correction of multiple hits to derive the estimated number of substitutions for each site, and 3) fit the site-specific substitution data to gamma distribution to obtain the shape parameter α This method is fast but its accuracy depends much on the accuracy of the estimated site-specific number of substitutions. The existing method has three shortcomings. First, it uses Poisson correction which is inadequate for almost any nucleotide sequences. Second, it does independent estimation for the number of substitutions at each site without making use of information at all sites. Third, the program implementing the method has never been made publically available. I have implemented in DAMBE software a new method based on the F84 substitution model with simultaneous estimation that uses information from all sites in estimating the number of substitutions at each site. DAMBE is freely available at available at http://dambe.bio.uottawa.ca

## 1 Introduction

Rate heterogeneity over sites arises in many different ways. For protein-coding genes, the substitution rate at the three codon positions, designated by r_1_, r_2_ and r_3_, respectively, are typically in the order of r_3_ > r_1_ > r_2_. This rate heterogeneity over codon positions has been attributed to two factors [1]. First, substitutions are always nonsynonymous at the second codon position, mostly nonsynonymous at the first codon positions, and frequently synonymous at the third codon position. Second, nonsynonymous substitutions at the second codon position involve very different amino acid, whereas those at the third codon position involve similar amino acids. This rate heterogeneity over codon sites depends on the intensity of purifying selection. For functionally unimportant genes, with pseudogenes as an extreme case, the difference among r_1_, r_2_ and r_3_ is small. For functionally important genes such as ribosome protein genes or histone genes, the difference becomes dramatic.

There are rate heterogeneity even among synonymous substitutions, mediated by differential tRNA availability [2, Chapter 9]. For example, in highly expressed yeast (*Saccharomyces cerevisize*) genes compiled in EMBOSS (Rice et al., 2000), we observe 314 AGA codons and only a single AGG codon. Both codons encode amino acid Arg. The difference in their usage can be attributed mainly to differential tRNA availability. The yeast genome encodes 11 tRNA^Arg/UCU^ genes for decoding AGA codons but only a single tRNA^Arg/CCU^ gene for decoding AGG without wobble-pairing (which is error-prone). Thus, tRNA-mediated selection would favor G→A replacement at the third codon position at AGR codons (where R stands for A or G), but not at other codons, e.g., AAR for Lys.

Protein genes have functionally important and unimportant domains that evolve at different rates. Some proteins (e.g., human insulin) are processed in such a way that some part of the polypeptides are discarded and do not appear in the mature protein, with those discarded segments having high evolutionary rates than those retained in the mature protein. Such rate heterogeneity among functional domains can often be detected by measuring the Ka/Ks ratio (nonsynonymous substitutions over synonymous substitutions) over a sliding window [3].

RNA genes such as rRNA and tRNA genes need to maintain a stable secondary structure for their normal function. The stem regions are typically conserved and the loop regions evolve rapidly [4]. The 18S rRNA has highly conserved and variable domains with both indel and nucleotide substitution events occurring predominantly in the eight variable domains [5].

## 2 Gamma distribution and gamma distances

Empirical studies of nucleotide substitution patterns [6-9] suggest that rate heterogeneity over sites is well approximated by gamma distribution defined as

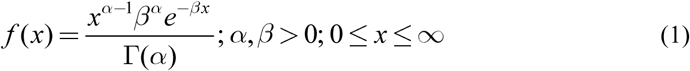

where α determines the shape of the gamma distribution and β is a scaling parameter. Gamma distances have been derived from commonly used substitution models such as JC69, K80, F84 and TN93. For JC69 [10-12]:

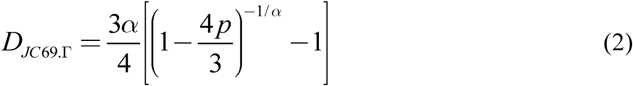

where p is the proportion of sites differing between the two alignment sequences from which D is desired.

The gamma distance for K80 [13, 14] is:

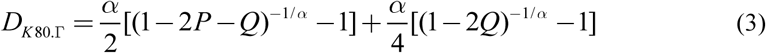

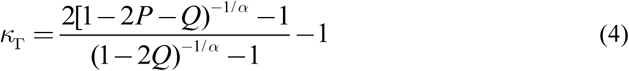

where P and Q are proportions of sites differing by a transition and a transversion, respectively, between two aligned sequences from which a distance is desired.

Contrasting the gamma distances with the regular non-gamma distances, Yang [15] noted the regularity that the gamma distance can be obtained by replacing ln(y) in the regular distance by –α(y^−1/α^ - 1) to obtain the gamma distance. Thus, to obtain the gamma distance for F84 [8], we can write down *at* and *bt* needed for the regular D_F84_ and use y_1_ and y_2_ to represent the arguments of the logarithms:

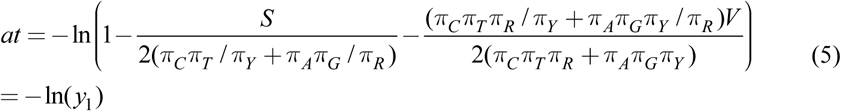

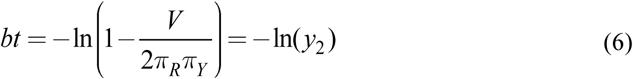

For D_F84.Γ_, we will express *at* and *bt* in the format of –α(y^−1/α^ - 1):

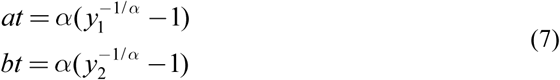

The equation for D_F84.Γ_ is of the same form as the regular D_F84_:

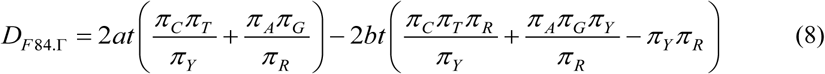

The gamma distance for TN93 [6] can be obtained the same way:

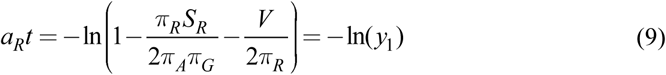

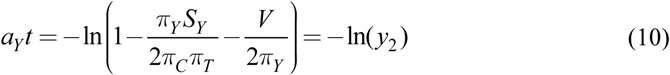

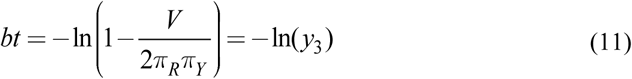

To obtain D_TN93.Γ_, we re-write

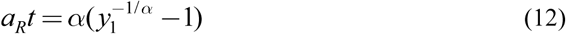

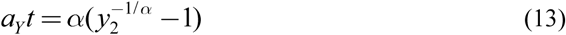

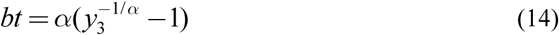

D_TN93.Γ_ is of the same form as for regular D_TN93_:

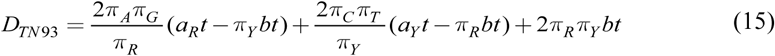

These distances have been implemented in DAMBE [16, 17] together with distance-based phylogenetic methods.

## 3 Estimating the shape parameter of a gamma distribution

We need to estimate α from empirical substitution data in order to compute the gamma distances in the previous section. The method by Gu and Zhang [9] uses the following probability density function [18] to estimate α:

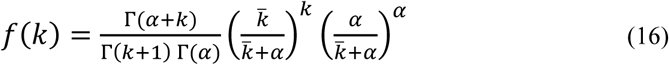

where k, instead of being integers, is replaced by the estimated number of substitutions per site, and, 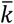 is mean k. Eq. (16) differs from that in Gu and Zhang [9] in that they wrote Γ(k+1) as ‘k!’ which is confusing because k is typically not an integer. Another minor difference is that they used D for 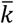 The advantage of using the probability density function in Eq. (16) over that in Eq. (1) is that it naturally use the number of monomorphic sites with k = 0, whereas the probability density in Eq. (1) is 0 for such sites.

The method is fast, but its accuracy depends on the accuracy of estimated k. The original method estimates k from a set of aligned sequences in two steps: 1) construct a phylogenetic tree from the aligned sequences and reconstruct ancestral sequences at internal nodes of the tree, and 2) perform pairwise comparisons along the tree between two nodes on each side of a branch to obtain observed number of substitutions per site, and apply correction for multiple hits to get the estimated number of substitutions per site. I illustrate these two steps below and highlight the two improvements in the last step as implemented in DAMBE [16, 17] in version 7.

### 3.1 Constructing a tree and infer ancestral sequences at internal node

Suppose we use the aligned sequences in the file VertCOI.fas (Fig. 1a) that comes with DAMBE installation [16, 17]. The file contains mitochondrial COI sequences from eight vertebrate species, with 1512 aligned sites and no indels. We may use either distance-based, maximum parsimony or maximum likelihood method to construct a phylogenetic tree. For this particular set of sequences, all these methods generate the same topology in Fig. 1b.

**Fig. 1.**
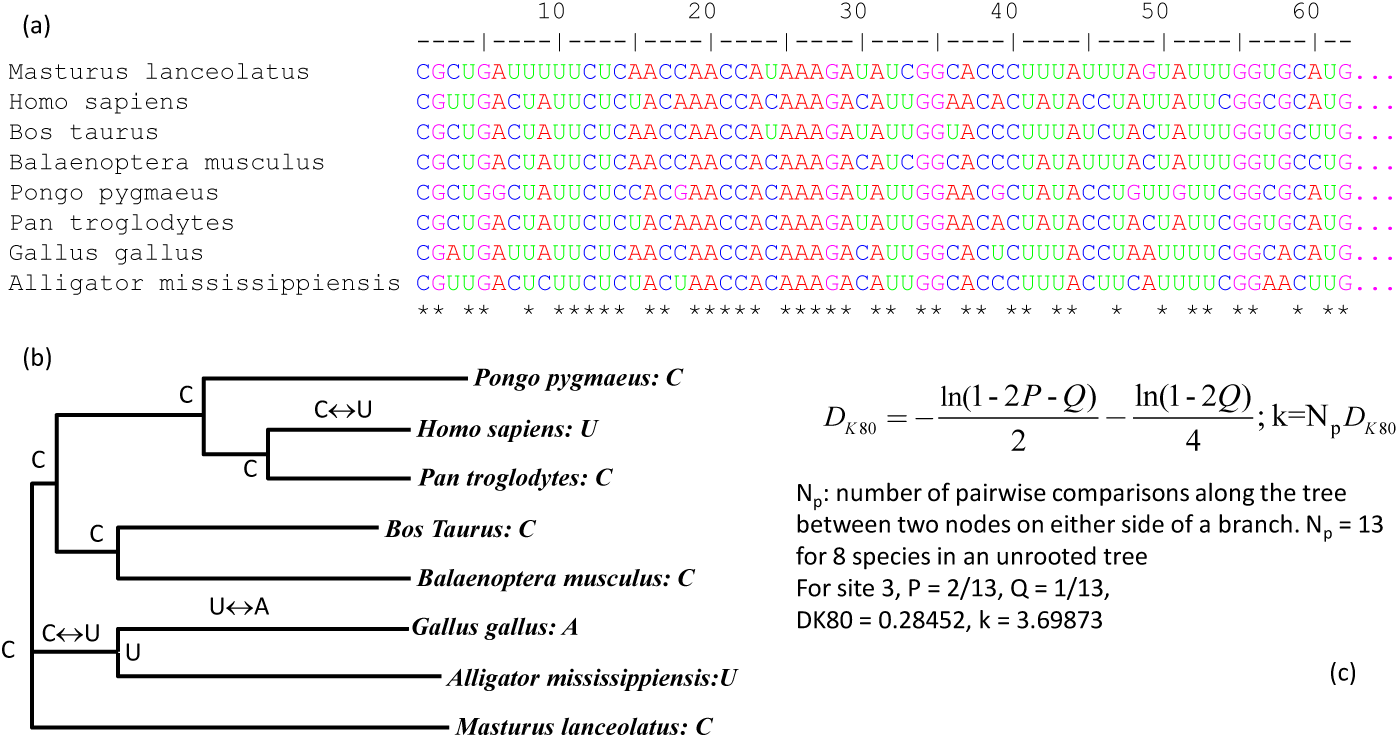
Phylogenetic reconstruction and estimation of site-specific number of substitutions. (a) Aligned mitochondrial COI sequences from eight vertebrate species (shown partially) in file VertCOI.fas that comes with DAMBE installation [16, 17]. (b) Phylogenetic tree with reconstructed ancestral states for site 3 in the aligned sequences. (c) Correcting for multiple hits by using K80 distance. k (= DK80*13 = 3.69873) is the estimated number of substitutions at site 3, where 13 is the number of branches in the tree.

Reconstructing ancestral states started with the application of the maximum parsimony algorithm [19, 20]. The approach was popularized by MacClade [21] and PAUP [22, 23] which included many options for handling non-uniqueness of reconstructed states. More recent approaches include empirical Bayes for nucleotide and amino acid sequences [24] and for amino acid sequences with options to incorporate structural information [25]. A web server taking the empirical Bayes approach is available [26] for reconstructing ancestral nucleotide, amino acid and codon sequences. The empirical Bayes approach assumes that the tree and branch lengths are correctly estimated. Because the tree and branch lengths in reality are uncertain, a more general hierarchical Bayesian approach [27, 28] has been investigated by using Markov chain Monte Carlo techniques for sampling phylogenetic trees and associated parameters. However, the reconstructed states from this general approach is similar to those from empirical Bayes except with greater uncertainty. DAMBE uses the same code in PAML [29] implementing the empirical Bayes approach for ancestral reconstruction. The reconstructed states for site 3 of the aligned sequences in Fig. 1a are shown next to the internal node of the tree in Fig. 1b. The first and second sites are monomorphic (Fig. 1a), so all internal node state is C for site 1 and G for site 2,

### 3.2 Perform pairwise comparisons along the tree between nodes on each side of a branch

For the first two sites, the number of substitutions is 0. For site 3, three substitutions were “observed” (assuming the reconstructed states for internal nodes), with two C⟷U transitions and one U⟷A transversion (Fig. 1b). Thus, the observed number of substitutions is three for site 3. Correcting for multiple hits by using the K80 model (Fig. 1c) is based on the following expected proportion of transitions and transversions are

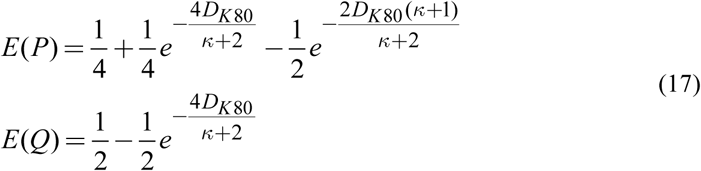

where D_K80_ is K80 distance and κ is the ratio of transition rate over transversion rate. Note κ is not to be confused with k in Eq. (16) which is the estimated number of substitution for a site. Replacing E(P) and E(Q) by the observed P and Q allow us to solve for D and κ (shown in Fig. 1c). For site 3, the observed P and Q are 2/13 and 1/13, respectively (Fig. 1b,c), which leads to D_K80_ = 0.28452. Note that this D_K80_ is the average number of substitutions per branch so, with 13 branches, k = DK80*13 = 3.6987.

The κ value, when estimated from one site, is naturally site-specific. Another site with a different substitution pattern will give us a different κ. It makes little sense that two neighboring sites should have different κ. More likely the same evolutionary process operate on all sites whose evolution should share the same κ. In addition, the estimation of D_K80_ above uses only one site of information (termed independent estimation or IE) which gives rise to many problems [30]. One of the problems involve inapplicable cases when D_K80_ cannot be estimated, especially when there are only eight or fewer sequences. Such inapplicable cases increase with more realistic substitution models, e.g., F84 or TN93. I illustrate the inapplicable cases below, and how such this problem can be avoided, or at least much alleviated, by simultaneous estimation (SE) using information from all sites.

Suppose we have a tree with 100 branches (N = 100), and we focus on three sites, S1, S2 and S3. The observed number of branches with transitional and transversional differences are shown in Table 1. For IE estimation, one simply substitute E(P) and E(Q) in Eq. (17) by the observed P and Q (which equal N_s_/N and N_v_/N, respectively) and then solve for D_K80_ and κ. You will encounter the frustration that you cannot estimate D_K80_ for S2 (Table 1).

**Table 1.**
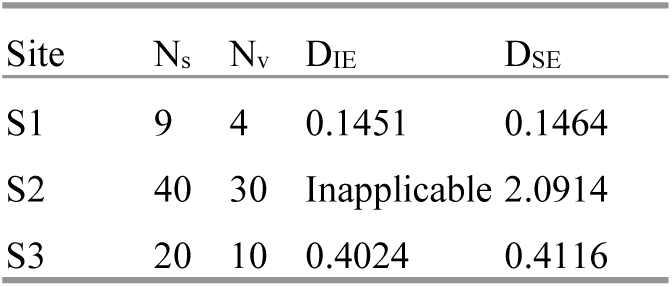
Observed number of transitional and transversional differences (Ns and Nv, respectively) for three sites (S1, S2 and S3) in a tree with 100 branches. Independently and simultaneously estimated K80 distances (DIE and DSE, respectively) are included. K80 distance cannot be computed for S2 using independent estimation method (labelled as ‘Inapplicable’).

| Site | $N_s$ | $N_v$ | $D_{IE}$ | $D_{SE}$ |
| --- | --- | --- | --- | --- |
| S1 | 9 | 4 | 0.1451 | 0.1464 |
| S2 | 40 | 30 | Inapplicable | 2.0914 |
| S3 | 20 | 10 | 0.4024 | 0.4116 |

For the SE method, we combine information from all sites to perform simultaneous estimation. We re-write Eq. (17) slightly differently to indicate that each site i has its site-specific D_i_ but all share the same κ:

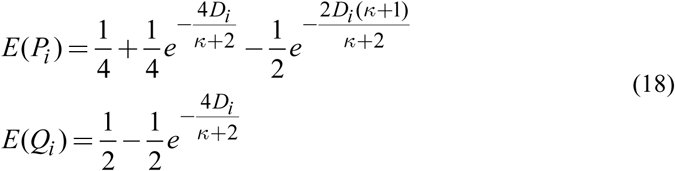

Now we can estimate all D_i_ and κ by maximizing the following log-likelihood:

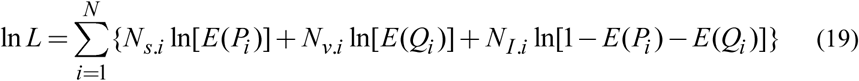

where N is the length of the aligned sequences, N_s.i_, N_v.i_ and N_I.i_ are recorded number of transitional difference, transversional difference and no difference from pairwise comparisons along the tree between nodes on each side of each branch. For example, N_s.3_ = 2, N_v.3_ = 1, and N_I.3_ = 10 (Fig. 1b). This simultaneous estimation is implemented in version 7 of DAMBE [16, 17] with the F84 substitution model.

### 3.3 Estimate the shape parameter (α)

Of the 1512 aligned sites in the VertCOI.fas file, 869 sites are monomorphic (k = 0). On the other extreme, one site recorded 5 transitions and 2 transversions, with k = 11.5642 (Table 2). The log-likelihood function for estimating α is

**Table 2.** Estimated k and the number sites with the same k value (N). f(k) is defined in Eq. (16) with values for α = 0.4977

| k | N | f(k) | N*ln[f(k)] | K | N | f(k) | N*ln[f(k)] |
| --- | --- | --- | --- | --- | --- | --- | --- |
| 0 | 869 | 0.5494 | -520.5370 | 4.7909 | 3 | 0.0870 | -7.3256 |
| 1.0429 | 121 | 0.1889 | -201.6587 | 5.1005 | 10 | 0.0256 | -36.6657 |
| 1.0962 | 60 | 0.1863 | -100.8256 | 5.4592 | 34 | 0.0409 | -108.7124 |
| 2.1799 | 107 | 0.1075 | -238.5962 | 5.8814 | 15 | 0.0732 | -39.2279 |
| 2.2996 | 59 | 0.1136 | -128.3390 | 6.3893 | 2 | 0.0256 | -7.3332 |
| 2.4340 | 50 | 0.1217 | -105.3292 | 7.2630 | 2 | 0.0143 | -8.4925 |
| 3.4244 | 50 | 0.0825 | -124.7452 | 7.8998 | 1 | 0.0414 | -3.1835 |
| 3.6278 | 41 | 0.0992 | -94.7368 | 8.6969 | 4 | 0.0211 | -15.4428 |
| 3.8594 | 64 | 0.1240 | -133.5730 | 10.2276 | 1 | 0.0042 | -5.4681 |
| 4.1260 | 18 | 0.0408 | -57.5806 | 11.5642 | 1 | 0.0058 | -5.1513 |

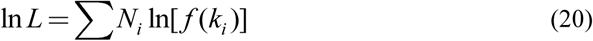

where f(k) is defined in Eq. (16). For any given α value, we can obtain the column of f(k) value according to Eq. (16). Summing up the N*ln[f(k)] column in Table 2 yields lnL. The largest lnL is -1942.924472, arrived when the α value equal to 0.4977 which is therefore the maximum likelihood estimate of α.

When α ≤ 1, the distribution of k is L-shaped (Fig. 2) with a strong right skew. Such a plot sometimes help us appreciate the rate heterogeneity over sites, i.e., some sites hardly change but some other site could have experienced nearly 12 substitutions (Table 2).

**Fig. 2.**
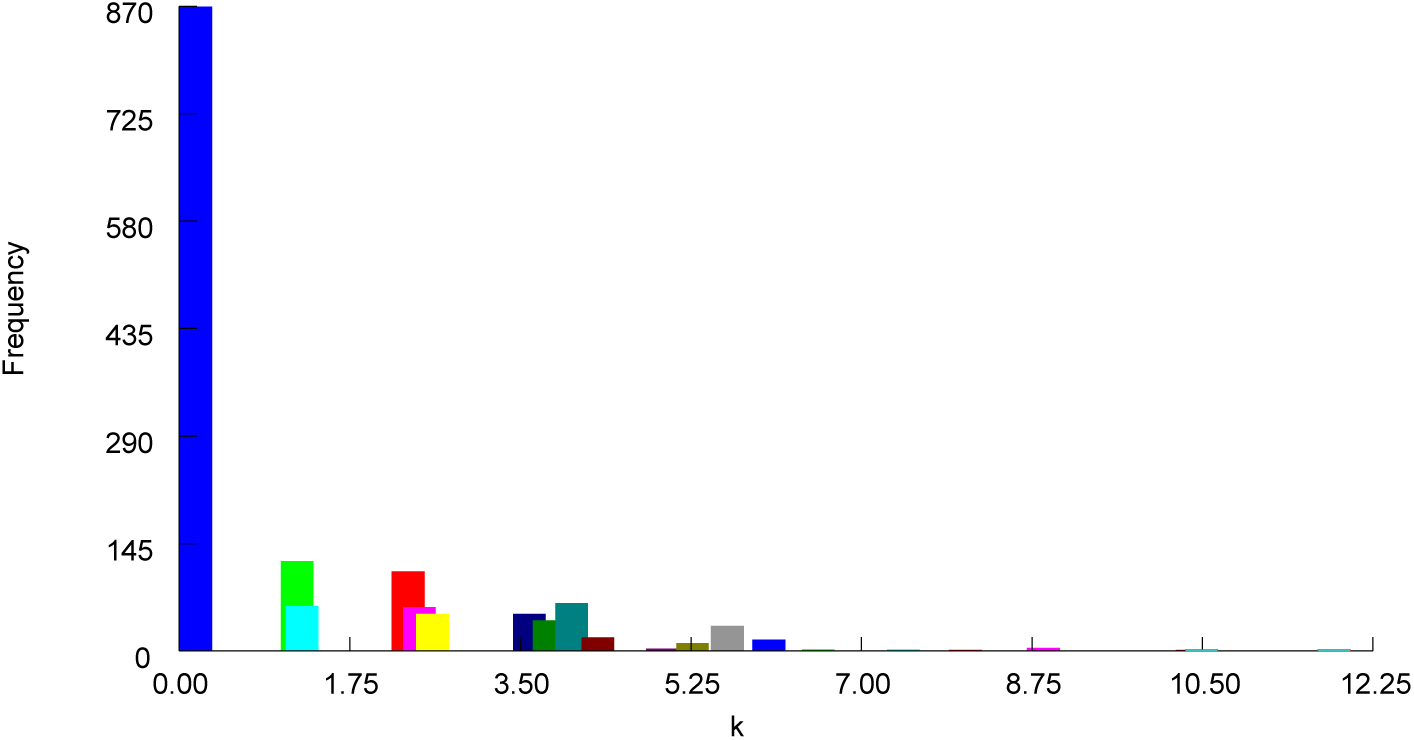
Frequency distribution of k (estimated site-specific number of substitutions). Output from DAMBE [16, 17].

## 4 Contrasting rate heterogeneity over sites among three codon positions

The three codon positions are expected to exhibit different heterogeneity [1]. Substitutions at the third codon positions are mostly synonymous and expected to occur at similar rates approximating neutral substitutions [30, 31]. Consequently, there should be relatively little rate heterogeneity at the third codon position. Substitutions at the first and second codon positions are almost all nonsynonymous. While substitutions at some functionally unimportant site might be close to neutral ones, some sites are expected to be strongly conserved. Thus, their rate differences among sites would span from nearly neutral rate at unimportant sites to almost no substitution at functionally constrained sites. This implies greater rate heterogeneity at the first and second codon positions than at the third codon position. This expectation is substantiated by empirical data, illustrated here with the aligned mitochondrial COI sequences from eight vertebrate species in the VertCOI.fas file that comes with DAMBE (Fig. 3). The rate heterogeneity is small at the third codon position (Fig. 3C) relative to those at the first and second codon positions (Fig. 3A,B). The third codon position also evolve in a more clock-like manner than the first and second codon positions [30, 31]. The α value is the smallest for the 2^nd^ codon position (Fig. 3B), which may be attributable to the observation that nonsynonymous substitutions at the 2^nd^ codon position involve substantially more different amino acids that at the 1^st^ codon position [1].

**Fig. 3.**
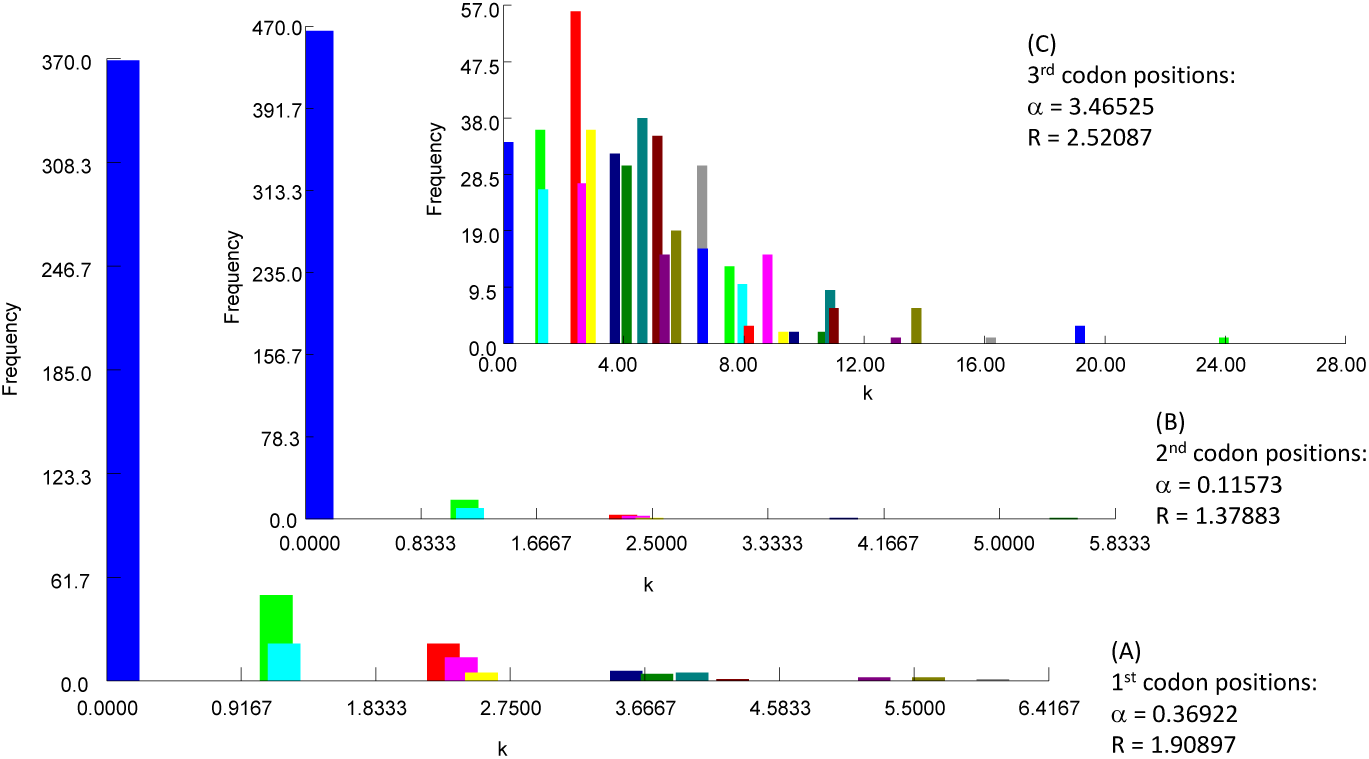
Contrast of rate heterogeneity among three codon positions of the vertebrate mitochondrial COI gene, together with estimated shape parameter α and transition/transversion ratio (R) for the F84 model used in DNAML. Output from DAMBE [16, 17].

## 5 Software availability

The most recent version of DAMBE is available free at http://dambe.bio.uottawa.ca. To access the function of estimating the shape parameter α, first read in a set of aligned sequences by clicking ‘File|Open standard sequence file’. Now click ‘Seq.Analysis|Substitution rate over site|Fit gamma distribution’.

## 6 Acknowledgement

This study is funded by the Discovery Grant from Natural Science and Engineering Research Council (NSERC, RGPIN/261252) of Canada.

